# Computationally prioritized drugs inhibit SARS-CoV-2 infection and syncytia formation

**DOI:** 10.1101/2021.04.15.440004

**Authors:** Angela Serra, Michele Fratello, Antonio Federico, Ravi Ojha, Riccardo Provenzani, Ervin Tasnadi, Luca Cattelani, Giusy del Giudice, Pia Anneli Sofia Kinaret, Laura Aliisa Saarimäki, Alisa Pavel, Vincenzo Cerullo, Olli Vapalahti, Peter Horvarth, Antonio Di Lieto, Jari Yli-Kauhaluoma, Giuseppe Balistreri, Dario Greco

**Affiliations:** Faculty of Medicine and Health Technology, Tampere University, Tampere, Finland; BioMediTech Institute, Tampere University, Tampere, Finland; Finnish Hub for Development and Validation of Integrated Approaches (FHAIVE), Tampere, Finland; Department of Virology, Faculty of Medicine, University of Helsinki, Helsinki, Finland; Drug Research Program, Faculty of Pharmacy, University of Helsinki, Helsinki, Finland; Synthetic and Systems Biology Unit, Biological Research Centre, Eotvos Lorand Research Network, Szeged, Hungary; Institute of Biotechnology, University of Helsinki, Helsinki, Finland; Department of Veterinary Biosciences, University of Helsinki, Helsinki, Finland; Department of Virology, University of Helsinki and Helsinki University Hospital, Helsinki, Finland; Institute for Molecular Medicine Finland, University of Helsinki, Helsinki, Finland; Department of Forensic Psychiatry, Aarhus University, Aarhus, Denmark; Queensland Brain Institute, The University of Queensland, Brisbane, Australia

**Keywords:** COVID-19, SARS-CoV-2, drug repositioning, drug design, virtual screening, 7-hydroxystaurosporine, bafetinib, syncytia, kinase inhibitors

## Abstract

New affordable therapeutic protocols for COVID-19 are urgently needed despite the increasing number of effective vaccines and monoclonal antibodies. To this end, there is increasing attention towards computational methods for drug repositioning and *de novo* drug design.

Here, we systematically integrated multiple data-driven computational approaches to perform virtual screening and prioritize candidate drugs for the treatment of COVID-19. From the set of prioritized drugs, we selected a subset of representative candidates to test in human cells. Two compounds, 7-hydroxystaurosporine and bafetinib, showed synergistic antiviral effects in our *in vitro* experiments, and strongly inhibited viral-induced syncytia formation. Moreover, since existing drug repositioning methods provide limited usable information for *de novo* drug design, we extracted and prioritized the chemical substructures of the identified drugs, providing a chemical vocabulary that may help to design new effective drugs.

## Introduction

The rapid diffusion of the COVID-19 pandemic has called for a prompt reaction from the biomedical research community. Although new vaccines have been developed as preventive options against the infection spreading (Deb et al., 2021; Thanh Le et al., 2020), little is still known about their efficacy (Callaway and Mallapaty; Mallapaty and Callaway), especially against variants of the virus. Furthermore, even if monoclonal antibody-based therapies represent an appealing option to treat the most severe cases of COVID-19, they are expensive and not easy to mass produce (Jaworski, 2020). Thus, more effective and affordable treatments for COVID-19 are still required to support medical intervention for the disease worldwide.

Currently available therapeutic options target components of the viral life cycle (e.g. nucleotide analogs or broad-spectrum antiviral drugs), the host immunological response (anti-inflammatory drugs or monoclonal antibodies), or vascular acute damage (antihypertensive and anticoagulant drugs) (Boccia et al., 2020; Giannis et al., 2020; Kaddoura et al., 2020; Pavel et al., 2021; Pujari et al., 2020; Yan et al., 2020). The angiotensin I converting enzyme 2 (ACE2) receptor and the transmembrane protease, serine 2 (TMPRSS2) may serve as therapeutic targets due to their crucial role in the initial phases of the viral infection (Hoffmann et al., 2020; Ni et al., 2020). Antivirals, especially antiretrovirals, represent the class of therapeutic agents which has been investigated to a larger extent (Frediansyah et al., 2021; Jomah et al., 2020). Various other drug classes were also proposed, such as anticancer drugs (e.g. kinase inhibitors) and antimicrobials (Lima et al., 2020; Malek et al., 2020). However, the majority of clinical trials focus on hydroxychloroquine alone or in combination with other compounds (38.65%), followed by immunotherapeutic (33.13%), and antiviral agents (9.20%) (Babaei et al., 2020).

Given the cost and time required for *de novo* drug development, drug repositioning is emerging as a viable solution (Dotolo et al., 2021; Jia et al., 2020; Napolitano et al., 2013; Rameshrad et al., 2020). Several efforts have been made to experimentally screen large libraries of candidate compounds (Ellinger et al., 2020; Gordon et al., 2020; Heiser et al., 2020; Touret et al., 2020). However, high-throughput screenings have a substantial cost in terms of time and resources. An *in silico* selection of candidate drugs with predicted desired activity would then reduce the need of experimental screenings. Several computational approaches to drug repositioning are available, either focusing on the targeting patterns of drugs to cellular components (Gordon et al., 2020), or based on the investigation of the mechanism of action (MOA) of drugs by omics profiling (Altay et al., 2020; Lima et al., 2020). However, there is a lack of integrative approaches that combine and enhance robustness of the predictions as well as provide useful information for *de novo* design of new compounds. Here, we propose a novel integrated computational approach to prioritize candidate drugs for treating COVID-19.

We prioritized compounds from the whole DrugBank library by virtual screening, and selected 23 candidates for *in vitro* infection assays to reveal drugs with potential antiviral activity against SARS-CoV-2. 7-Hydroxystaurosporine and bafetinib showed significant antiviral effect in human epithelial cells. Interestingly, the two drugs also revealed a synergistic effect in blocking virus-induced syncytia formation. Furthermore, our analytical framework allows the prioritization of relevant chemical substructures, which could serve as a molecular fragment library for *de novo* drug development for COVID-19.

## Results and discussion

### Computationally aided drug prioritization identifies a subset of candidate drugs for COVID-19

We focused on multiple aspects relevant for the prioritization of candidate drugs for COVID-19 treatment (Figure 1A). We analyzed publicly available transcriptomics data to characterize both the MOA of drugs (Igarashi et al., 2015) as well as the disease-specific alterations (Blanco-Melo et al., 2020; Daamen et al., 2021). We highlighted chemical substructures present in drugs that are able to: (1) alter the expression of genes related to the initial phases of the viral infection (Gordon et al., 2020), (2) revert the virus-induced transcriptomic alteration, and (3) target key genes of the host response to the SARS-CoV-2. Moreover, we identified structural properties of drugs predictive of the deregulation level of the ACE2 receptor, the transmembrane protease TMPRSS2, and the cell surface proteolytic enzymes cathepsin B (CTSB) and procathepsin L (CTSL). Finally, we retrieved chemical substructures from drugs that were identified as active in previous screenings for COVID-19.

**Figure 1:**
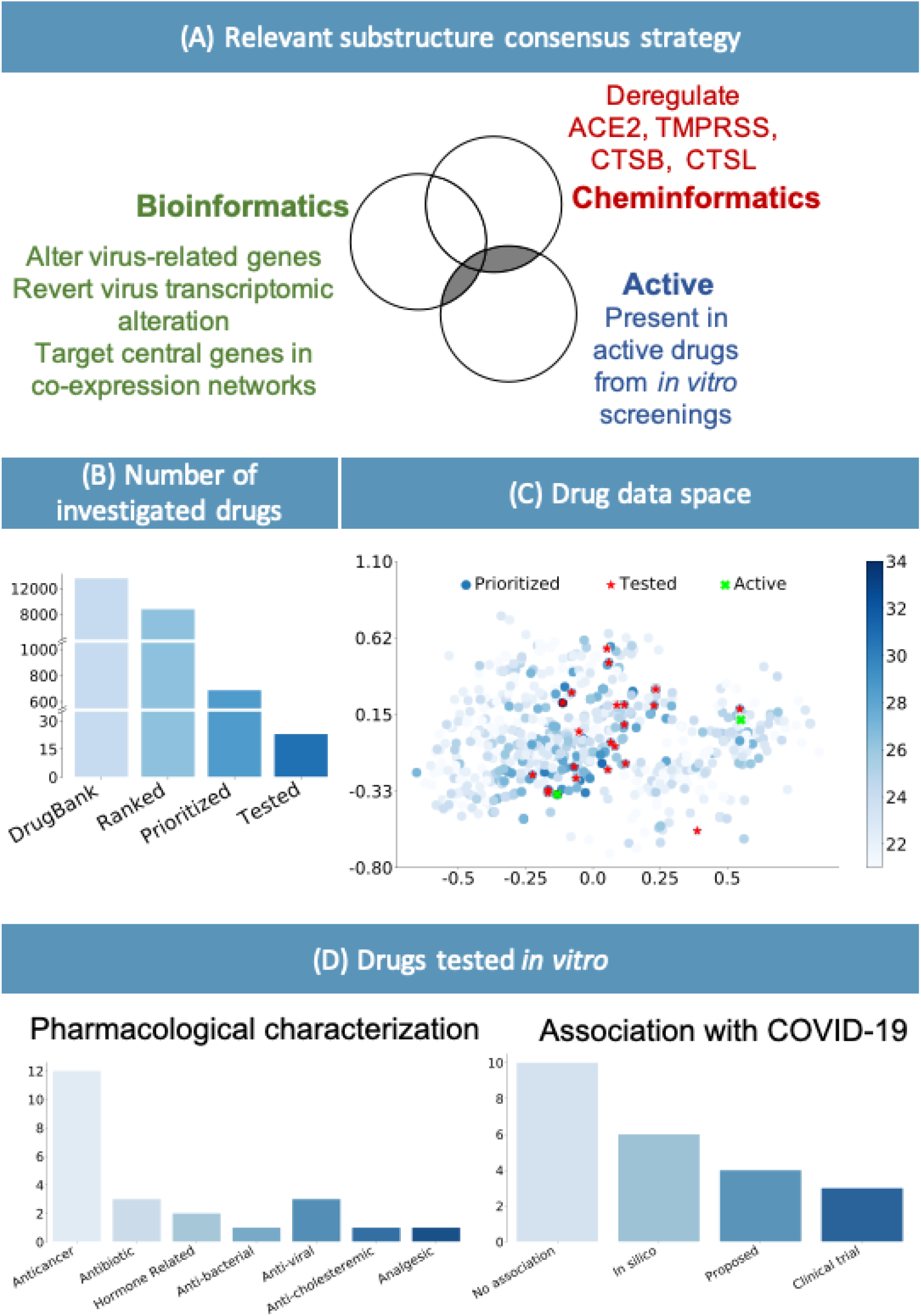
(A) Consensus strategy to identify relevant chemical substructure, using bioinformatics and cheminformatics methods as well as experimental results from published literature. (B) The suggested approach allows reducing the number of experimental tests: the whole DrugBank database was filtered to less than 2000 relevant drugs and *in vitro* testing was performed on 23 candidates. (C) Graphical representation of the prioritized drugs. The shade of blue represents the number of chemical substructures identified in (A), present in the drugs. The 23 selected compounds are shown in red. They were selected among the drugs sharing the most relevant substructure as well as satisfying practical logistic criteria. Of the 23 drugs, the 2 highlighted in green have been experimentally identified as active. (D) Pharmacological characterization and description of known association with COVID-19 of the 23 tested drugs. *In silico* refers to drugs derived from *in silico* studies, while proposed refers to drugs suggested for their potential therapeutic role in literature.

To identify candidate drugs effective against COVID-19, we computationally prioritized drugs from the DrugBank database (Wishart et al., 2006, 2018) with PubChem-available fingerprints (Figure 1B). We prioritize these drugs according to the presence of chemical substructures expected to interfere with disease-associated biological processes. From 8000 total drugs examined, we focused on the top 700 to select a set of relevant candidates for further experimental validation. Our inclusion criteria is a trade-off between the selection of a set of drugs that best represents the top of the prioritized list, based on their chemical substructures (Figure 1C), and a number of practical considerations such as price, availability, shipping time, and ease of storage. Taking into account the aforementioned criteria, we validated our method by performing an *in vitro* biological evaluation of 23 selected drugs.

The majority of drugs currently in clinical trials for COVID-19 treatment present either antiviral or immunomodulating properties, with the aim of targeting the viral life cycle and alleviating the lung-damaging symptoms (Babaei et al., 2020; Wang and Guan). Kinases play an important role in multiple of these biological processes, and therefore, different kinase inhibitors were proposed for COVID-19 treatment (Saha et al., 2020; Weisberg et al., 2020). These compounds show pharmacodynamic properties allowing the dual goal of mitigating both host immunological response and antiviral activity (Weisberg et al., 2020). Twelve out of the 23 identified drugs relate to oncological treatments, and eight of them act as kinase inhibitors (Figure 1D). Similarly to antiviral drugs, anticancer drugs may, indeed, target biological processes which have a crucial role in modulating the organism immune response, cell division and death, cell signaling, and microenvironment generation (Borcherding et al., 2020). Several studies about repurposing of anticancer drugs to treat COVID-19 already exist (Borcherding et al., 2020; Braga et al., 2021; Saini et al., 2020, 2020). Roshewski et al. showed that acalabrutinib, a selective Bruton tyrosine kinase inhibitor, can mitigate the hyperinflammatory immune response characterising the most severe cases of COVID-19 (Roschewski et al., 2020). The histone methyltransferase inhibitor pinometostat, instead, seems to decrease the level of NFkB, one of the main players of the immunological response (Hirano and Murakami, 2020), and to mitigate the host-response against infections (Marcos-Villar and Nieto, 2019). Inhibitors of intracellular calcium homeostasis seem to block virus-induced cell–cell fusion (known as syncytia formation), one of the hallmarks of severe SARS-CoV-2 infection (Braga et al., 2021; Bussani et al., 2020).

Our analysis successfully highlighted several drugs that are either under investigation or acted effectively against SARS-CoV-2 infection. The selected 23 candidates for further investigation comprise anticancer, antimicrobial, and antiviral drugs (Figure 1D). Drugs from all three classes showed to lower the virus titre and to tune down the cytokine storm syndrome in the most severe cases of the disease (Ginsburg and Klugman, 2020). As expected, our approach identified antiviral drugs (against hepatitis C), which were already predicted as a COVID-19 treatment, or are currently in clinical trials against SARS-CoV-2 infection^1^ (Jockusch et al., 2020; Mevada et al., 2020). Among the three identified antibiotics, delafloxacin, a fluoroquinolone antimicrobial agent, was also studied for its antiviral activity at the early stages of the COVID-19 pandemic (Karampela and Dalamaga, 2020).

Altogether, approximately half of the tested candidate drugs were proposed as a potential COVID-19 treatment (Figure 1D). This demonstrates the capability of our methodology to identify potential drug candidates and to highlight a new set of existing compounds without previous association to COVID-19 or SARS-CoV-2.

### Validation in human cells confirmed 7-hydroxystaurosporine and bafetinib as potential COVID-19 treatment

To experimentally validate our computational predictions, we tested the set of candidate drugs at different concentrations (0.09 μM, 0.9 μM, and 9 μM) on HEK-293T cells stably expressing human *ACE2* and *TMPRSS2* (HEK-293T-AT) (Cantuti-Castelvetri et al., 2020). Cells were infected with WT SARS-CoV-2, at a multiplicity of infection (MOI) of 0.5 infectious units (i.u.) per cell for 16-18 h. The percentage of infected cells was determined by immunofluorescence detection of viral proteins followed by automated imaging and image analysis. Out of the 23 drugs, 7-hydroxystaurosporine and bafetinib, showed a statistically significant inhibition of the number of virus-infected cells when tested at 9 μM, showing relative infection values of 0.51 and 0.69, respectively (Figure 2A). Ponatinib, instead, significantly inhibited the infection but also displayed a strong cytotoxic effect (Figure 2A), and was therefore excluded from further analysis.

**Figure 2:**
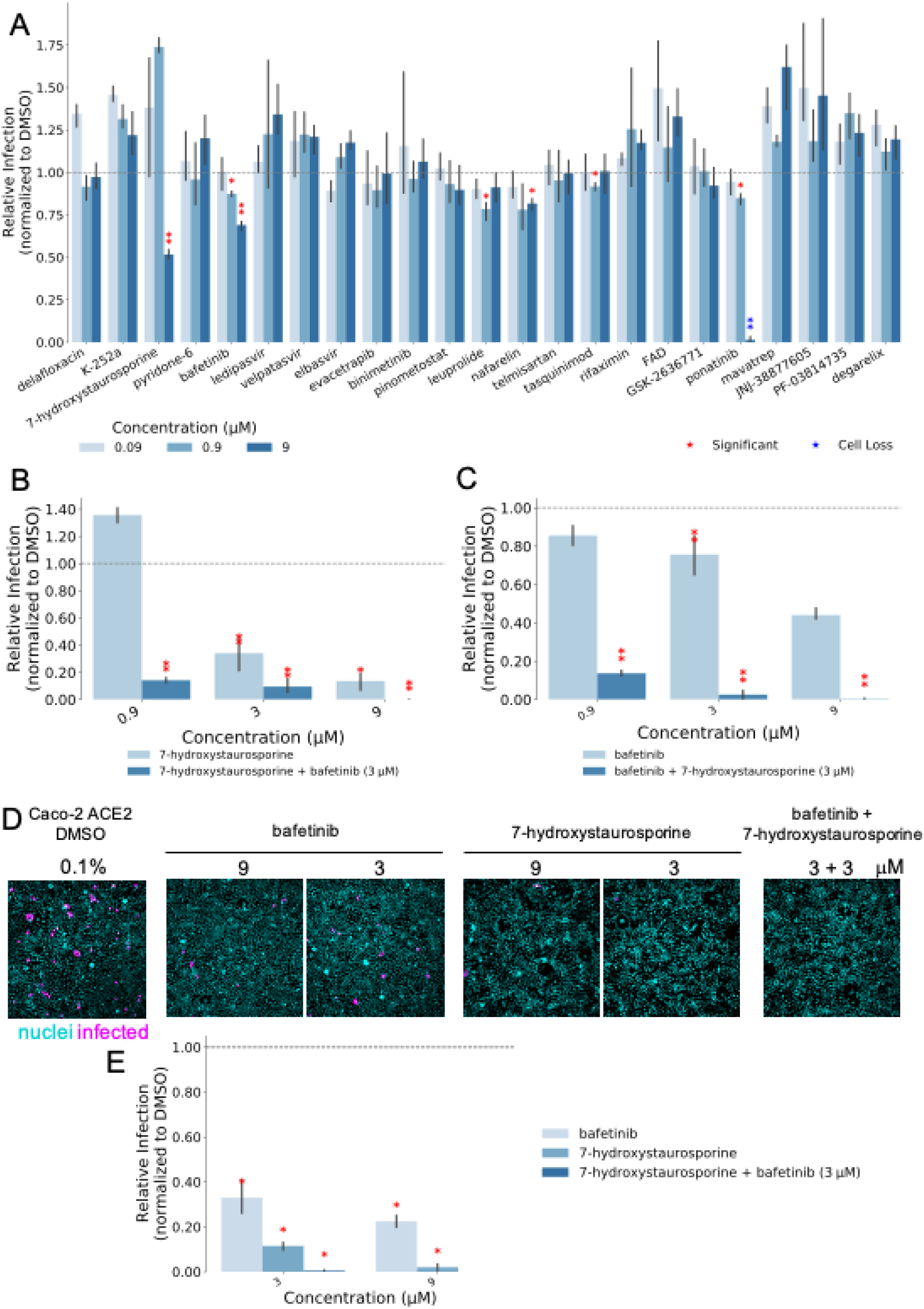
A) Percentage of infected cells after drug treatments normalized by the median of the dimethyl sulfoxide (DMSO) control. Each drug was added 45 min before infection and infected cells fixed 16 h later; Red asterisks show significant *p* values (<0.05) for the one-tailed *t*-test between each treatment and the DMSO. B) Combined effect of 7-hydroxystaurosporine and bafetinib added 2 hours before infection (hbi) at indicated concentrations. Cells fixed 16 hours post-infection (hpi). C) Combined effect of bafetinib and 7-hydroxystaurosporine added 2 hbi at indicated concentrations. Cells fixed 16 hpi. D) Representative fluorescence images of Caco2-ACE2 cells treated with indicated drugs 2 hbi. Cells fixed 16 hpi; blue=nuclei, magenta=infected cells. E) Quantification of experiment in D; values normalized to DMSO controls. All values represent the averages of 3 experiments. Error bars indicate the standard deviation.

7-Hydroxystaurosporine is an antineoplastic agent with potent *in vitro* and *in vivo* activities, and its capability of sensitising a variety of cell lines *in vitro* was previously described (Lara et al., 2005; Monks et al., 2000). 7-Hydroxystaurosporine is often used in combination with other drugs for its synergistic effect of enhancing cytotoxic effect in human cancer cells, e.g. in treating leukemia^2^ (Monks et al., 2000; Sampath et al., 2006; Senderowicz, 2001). To our knowledge, no connection between 7-hydroxystaurosporine and SARS-CoV-2 emerged in terms of possible COVID-19 treatment. Bafetinib, is a second generation tyrosine kinase inhibitor prescribed against Philadelphia chromosome–positive chronic myelogenous leukemia (Santos et al., 2010). In addition, bafetinib was recently identified as a SARS-CoV-2 inhibitor in other drug repurposing studies (Bouhaddou et al., 2020; Drayman et al., 2020). Bouhadduo et al. proposed and tested bafetinib as a possible COVID-19 drug, modifying the phosphoproteome of SARS-CoV-2 infected cells *in vitro* (Bouhaddou et al., 2020).

### 7-Hydroxystaurosporine and bafetinib inhibit SARS-CoV-2 infection

Next, we tested whether a combination of the two identified compounds would result in stronger antiviral activity without compromising cell viability. For this, we performed two additional infection assays in which we exposed the cells to 7-hydroxystaurosporine and bafetinib alone, or in combination. In the first assay, the concentration of bafetinib was fixed at 3 μM, while the concentration of 7-hydroxystaurosporine varied as 0.9 μM, 3 μM, and 9 μM (Figure 2B). In the second assay, the concentration of 7-hydroxystaurosporine was fixed at 3 μM, while the concentration of bafetinib varied as in the first assay (Figure 2C). The cells were exposed to the drugs 2 hours before infection (hbi).

These results indicate that 7-hydroxystaurosporine had a significant inhibitory effect on viral infection already at 3 μM (Figure 2B), without remarkable cytotoxic effects. The combination with bafetinib resulted in an increased antiviral activity, reducing the number of infected cells by more than 80%, without compromising cell viability (Figure 2B). The same applies for bafetinib, which showed a moderate but significant antiviral activity already at 3 μM and the activity was even stronger when combined with 7-hydroxystaurosporine (Figure 2C).

To test if the drugs blocked SARS-CoV-2 cell entry, which occurs during the first hour of infection (Koch et al., 2020), or a post-entry step of the virus life cycle, we added the drugs 2 hours post-infection (hpi) and quantified the fraction of infected cells 16 hpi. We observed a similar antiviral effect in cells treated after infection, indicating that the drugs inhibit mainly a post-entry step of infection.

The effect of the two drugs was tested in a different cell line, Caco-2 cells stably overexpressing *ACE2* (Caco2-ACE2) and endogenously expressing *TMPRSS2* (Figure 2D). The antiviral effect of 7-hydroxystaurosporine, bafetinib, and of the two drugs combined, was stronger compared to HEK-293T-AT. At a concentration of 3 μM, the infection inhibition was more than 60%, 80%, and 90%, respectively (Figures 2D–E).

Both bafetinib and 7-hydroxystaurosporine exhibited a concentration-dependent inhibition of viral infection (Figure 3) and predicted benchmark dose (BMD) values of 1.22 μM and 5.09 μM respectively (Figures 3A–B). The predicted BMD values of the combination were 0.63 μM and 0.65 μM (Figures 3C–D), suggesting that lower concentrations might be already effective to reduce the infection rate.

**Figure 3:**
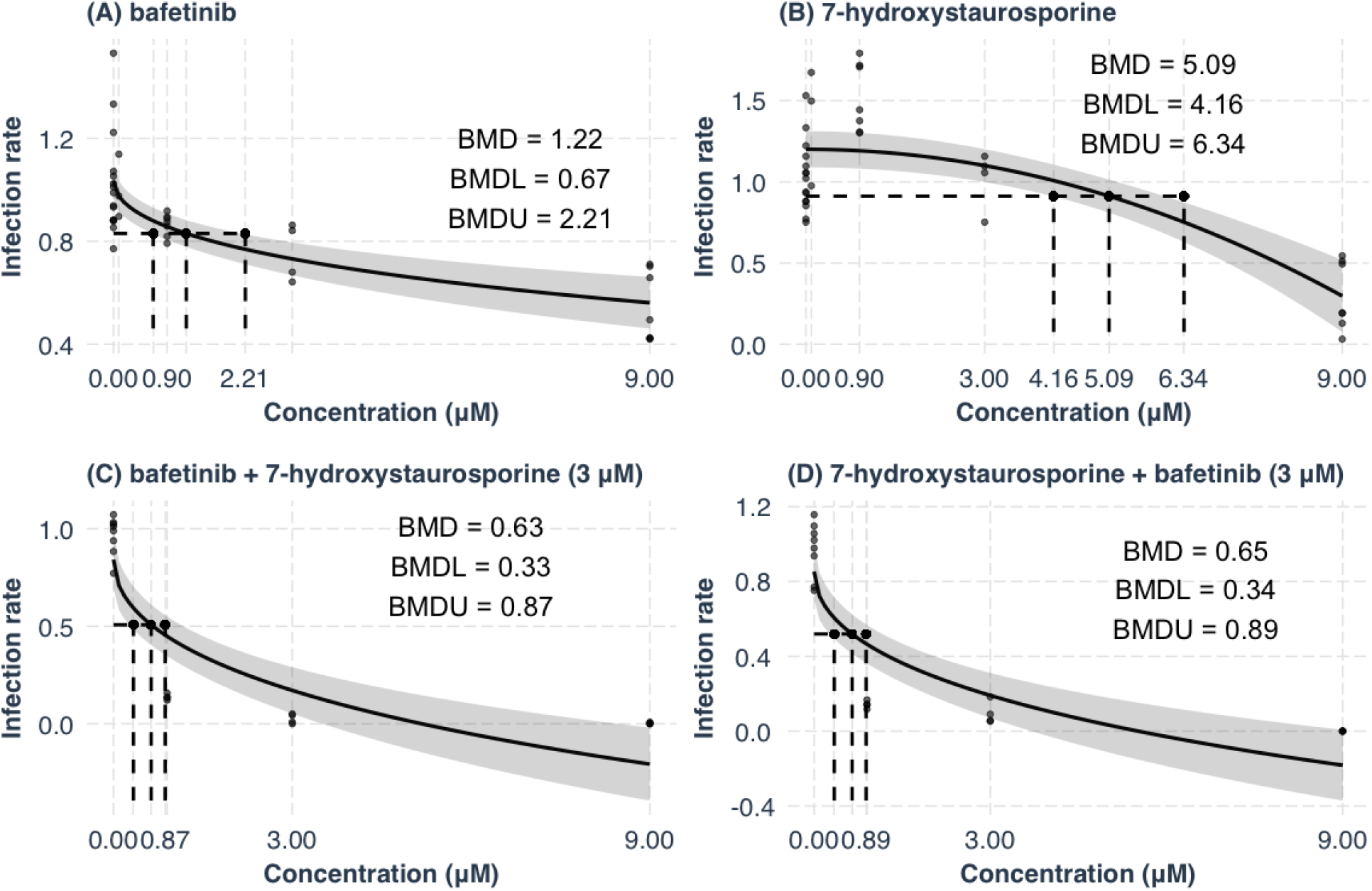
Concentration–response curve analysis. The benchmark dose–response (BMD) values and their lower (BMDL) and upper (BMDU) bounds were computed for bafetinib (A), 7-hydroxystaurosporine (B) and their combinations (C–D). The y-axes show the infection rate normalized by the one measured in the DMSO. Bafetinib and 7-hydroxystaurosporine were tested at 0.09 μM, 0.9 μM, 3 μM, and 9 μM, while in combination they were tested at 0.9 μM, 3 μM, and 9 μM. The BMD, BMDL, and BMDU in (C) refer to the experiments performed where 7-hydroxystaurosporine was combined with a fixed concentration (3 μM) while bafetinib concentration varied.

### 7-Hydroxystaurosporine and bafetinib synergistically block SARS-CoV-2–induced cell–cell fusion

The ability of some pathogenic human viruses, including SARS-CoV-2, to induce cell–cell fusion, a phenomenon known as syncytia formation (Buchrieser et al., 2021), was linked to viral spreading, pathogenicity and tissue damage *in vivo* (Bussani et al., 2020). When infected by SARS-CoV-2, HEK-293T-AT cells efficiently fuse, forming large multinucleated cells (Figure 4A). In addition to a reduction in the number of infected cells, a machine learning–assisted image analysis revealed that 7-hydroxystaurosporine, at 4.5 μM, strongly inhibited virus-induced cell–cell fusion, reducing the average nuclear content per cell by more than 80% (Figure 4A–B). The combination with bafetinib increased this inhibitory effect, reducing the formation of large multinucleated cells by more than 90% and 60%, at 4.5 μM and 2.25 μM, respectively (Figure 4B). At the lowest concentration, 7-hydroxystaurosporine had a moderate inhibitory effect on syncytia formation, whereas the combination with bafetinib significantly blocked cell–cell fusion even at 1.125 μM (Figure 4B).

**Figure 4:**
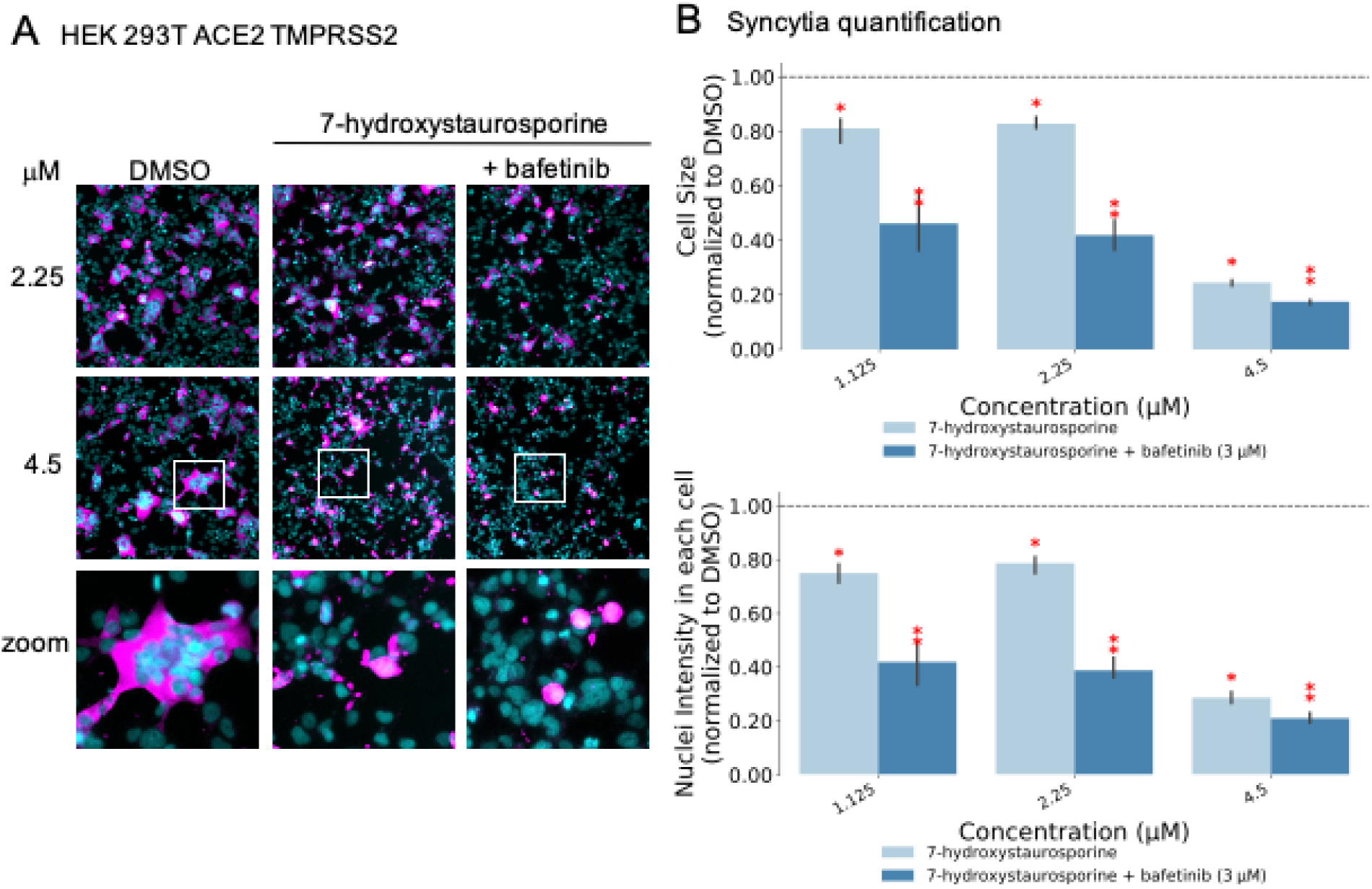
7-Hydroxystaurosporine and bafetinib inhibit virus-induced syncytia. A) Representative fluorescence images of HEK-293T-AT cells treated with indicated drugs 1 hour before infection. Cells fixed 16 hours post-infection; blue=nuclei, magenta=infected cells. Zoomed areas from each image are indicated by white boxes; B) Quantification of cell size and nuclear content from the experiment in A; values normalized to the median of DMSO controls. All values represent the averages of 3 experiments. Error bars indicate the standard deviation. Red asterisks show significant *p* values (<0.05) for the one-tailed *t*-test between each treatment and the DMSO.

By targeting ABCB1 and ABCG2 transporters, bafetinib is known to increase the intracellular accumulation of anticancer drugs by blocking the drug efflux (Zhang et al., 2016). This could be a plausible mechanism for how this inhibitor enhances the effect of 7-hydroxystaurosporine. Interestingly, in addition to allowing drug efflux, this group of ABC transporters was recognised for their role in syncytialization (Manceau et al., 2012). Buchrieser et al. and Ou et al. highlighted connections between multinucleated syncytial cells and, similarly to other viruses such as measles, respiratory syncytial virus (RSV), and MERS, SARS-CoV-2 induces cell–cell fusion, a phenomenon particularly evident in in severe COVID-19 (Buchrieser et al., 2020; Ou et al., 2020). Sisk et al. demonstrated that the membrane fusion is blocked in the presence of Abl kinase inhibitors (imatinib, GNF2 and GNF5), and thus, prevented syncytia formation in coronavirus (infectious bronchitis virus) spike protein–induced Vero cells (Sisk et al., 2018). More recently, another study demonstrated that the fusogenic activity of the viral spike can be inhibited by targeting cellular factors (Braga et al., 2021).

### Structural comparison between 7-hydroxystaurosporine and bafetinib

As 7-hydroxystaurosporine and bafetinib exhibited the highest inhibitory activity against SARS-CoV-2 infection at low micromolar concentrations, we investigated further their 2-D structures and 3-D geometries. In terms of molecular fingerprints, most of the common substructures highlighted aromaticity-related similarities. In fact, based on their 2-D structures (Figure 5A), both candidates possess a highly conjugated π-bond system with 22 and 27 sp^2^-hybridized atoms for 7-hydroxystaurosporine and bafetinib, respectively. Fused aromatic rings and high degree of conjugation suggest a planar geometry, which was confirmed when we generated and compared their 3-D models (Figures 5B–D). Despite the different nature of their 2-D structural scaffolds, the generated 3-D geometries overlayed and displayed a 67% shape-similarity, which may partially explain their antiviral activity in the biological assays. The 3-D conformation generated for bafetinib is in line with the model presented by Zhang et al. (Zhang et al., 2016), with the ligand docked into the binding pocket of ABCG2 efflux transporter. An inhibition of such efflux transporter also supports the possible explanation for the synergistic effect when the compounds were used in combination. However, as 7-hydroxystaurosporine and bafetinib lowered the infection rate also when tested alone, a clear explanation for their intrinsic antiviral activity remains yet unknown. Of note, the drug candidate K-252a shares the same aromatic and planar core of 7-hydroxystaurosporine, however, it did not show inhibitory activity in the biological assays. The lack of its intrinsic activity may be due to the missing hydroxy group in the isoindolinone moiety and/or to the different substituents (e.g. the lack of a basic functional group) and number of the sp^3^-hybridized carbon atoms in the conformationally distinct aliphatic heterocyclic moieties, i.e. tetrahydro-2*H*-pyran and tetrahydrofuran in 7-hydroxystaurosporine and K-252a, respectively.

**Figure 5.**
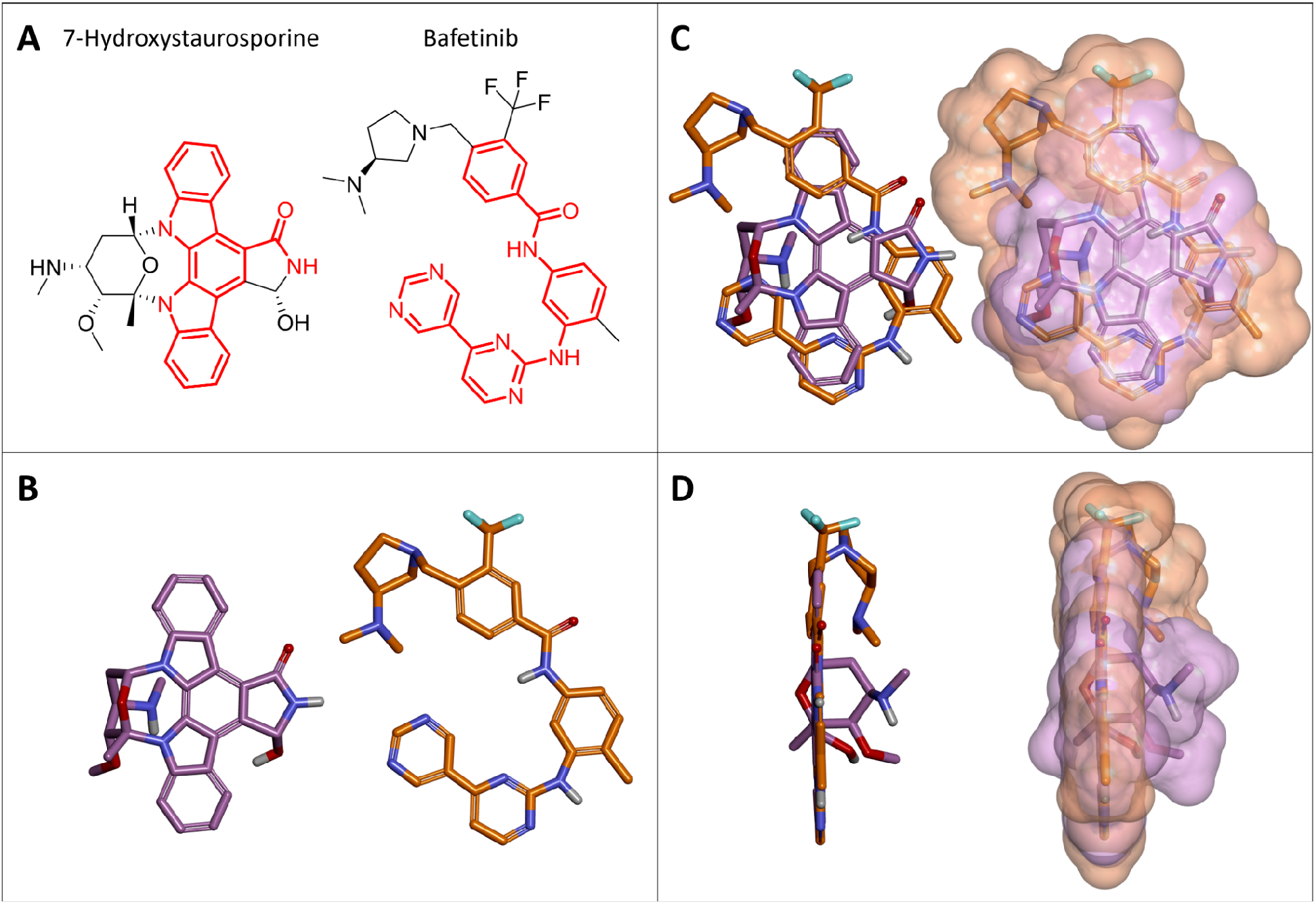
Structural comparison between 7-hydroxystaurosporine and bafetinib. (A) 2-D structures of 7-hydroxystaurosporine (left) and bafetinib (right); the conjugated π-bond system is highlighted in red. (B) 3-D structures of 7-hydroxystaurosporine (left) and bafetinib (right). (C) Front view and (D) side view of 7-hydroxystaurosporine and bafetinib 3-D structural overlay with (right) and without (left) solvent accessible surface. Color code for (B), (C), and (D): carbon atoms and solvent-accessible surfaces are shown in lilac and orange for 7-hydroxystaurosporine and bafetinib, respectively; oxygen atoms in red; nitrogen atoms in blue; fluorine atoms in cyan; and hydrogen atoms in white.

### Conclusive remarks

We prioritized the DrugBank database for potential SARS-CoV-2 inhibitors by using a computational procedure. To achieve that, we integrated established methodologies from the fields of bioinformatics and cheminformatics. We experimentally assayed 23 representatives of the prioritized library and found that two drugs, bafetinib and 7-hydroxystaurosporine, showed significant inhibition of viral infection. In addition, an image analysis of the infected versus treated cells showed that the formation of multinucleated syncytial cells was also significantly reduced. Unexpectedly, when combined, the two drugs exerted an even stronger, synergistic inhibition of viral infection as well as cell–cell fusion inhibition at lower concentrations. Further *in vitro* experimentation showed that the drugs in combination were still effective one hour after the infection of the cells, suggesting that they may hinder some post-entry mechanism of the virus. The prioritization methodology we developed allowed us to make an informed decision about which drugs to test, out of almost 8000 drugs from DrugBank, thus significantly reducing the costs of drug repurposing, while simultaneously increasing the success rate. Also, our results are not limited to candidate repositionable drugs, but include a characterization in terms of chemical substructures relevant to the viral system considered. Indeed, we conjecture that this set of chemical substructures can be further exploited in the context of scaffold-based *de novo* drug design. We believe that our integrated approach could significantly help the repurposing of drugs for other viral diseases, as well as other complex pathologies.

## Materials and Methods

### RNA-Seq data preprocessing

Human transcriptomics datasets analysed in this study were retrieved from the Gene Expression Omnibus (GEO) repository, annotated with the GEO ID GSE147507 (Blanco-Melo et al., 2020; Daamen et al., 2021). The datasets are composed as following: human lung biopsies of SARS-CoV-2 infected patients and uninfected control; A549 cell line infected with SARS-CoV-2, A549 cell line infected with SARS-CoV-2 overexpressing ACE2, Calu-3 cells infected with SARS-CoV-2; NHBE cell line infected with SARS-CoV-2. For each of the cell lines, the mock treated lines were collected to be used as controls for the expression analysis. The preprocessing of human transcriptomics datasets was carried out starting from the raw counts provided within the GEO record. Low read counts were filtered by applying the proportion test method implemented within the NOISeq Bioconductor package (Tarazona et al., 2011). Filtered counts were then normalized through the upper quartile method implemented in the NOISeq package. Differential expression analysis was carried out by using the DESeq2 Bioconductor package (Love et al., 2014) and the *p* values were adjusted using the Benjamini-Hochberg method (Benjamini and Hochberg, 1995).

### Co-expression network inference and analysis

Co-expression networks were inferred for both the human biopsies and all the infected cell lines by using the clr algorithm (Faith et al., 2007) implemented in the minet package (Meyer et al., 2008) with Pearson correlation as the estimator. The expression values of the differentially expressed genes were used to infer the networks. The genes in each of the networks were ranked based on several centrality measures, including degree, closeness, clustering coefficient, betweenness and eigenvector, in addition to a biological significance score calculated as follows: |*log*_2_ (*FC*) ×− *log*_2_ (*p* − *value*) | by using the INfORM tool (Marwah et al., 2018).

### Open TG-GATEs data preprocessing

Raw microarray data for 129 drugs were downloaded from the Open TG-GATE repository^3^ (Igarashi et al., 2015). Drugs were tested in *in vivo* and *in vitro* rats samples (both in liver and kidney) with three dose levels at four different time points. Samples were imported in R by using the justRMA function (Irizarry et al., 2003) from the R library Affy (Gautier et al., 2004). Outliers were identified by using the RLE, NUSE from the affyPLM package (2005) and the slope of the RNA degradation curve implemented in the affy package (Gautier et al., 2004). A sample was removed from the analysis, when marked as outlier by at least two out of the three methods. The probes were annotated to Ensembl genes (by using the rat2302rnensgcdf (v. 22.0.0) annotation file from the brainarray website^4^, and the resulting expression matrix was quantile normalized by using the normalizeQuantile function from the limma package. Eventually, only the subset of samples tested in *in vivo* rat liver tissue with a single exposure experimental setup were considered for further analysis and the ensembl genes were mapped to gene symbols by using the AnnotationDBi package (Pagès et al., 2018).

### Dynamic dose-responsive point of departures

For each drug of the Open TG-GATEs, the dynamic dose-dependent MOA and the corresponding point-of-departure (POD) were identified through the use of the TinderMIX software (Serra et al., 2020a). Starting from the pairwise log2-fold-change (log2FC) of each gene (computed as the difference between the log2 expression values of each pair of treated and control samples), first-, second- and third-order polynomial models were fitted. For the best-fitting model its contour plot was computed as an effect map. Next, the contour plot of each gene was evaluated to identify an area showing a dynamic dose-response, *i*.*e*. an area where the expression changes monotonically in respect to the dose once an activity threshold has been reached. As for the classical BMD analysis, an activity threshold of 10% was selected (Serra et al., 2020b; Thomas et al., 2007; Yang et al., 2007). If such an area was identified, the gene was considered to be altered in a dynamic dose-dependent manner.

Eventually, based on the lowest dose and earliest time of activation, the genes were labelled with an activation label that specifies its POD. The time-dose effect map was divided into a 3 by 3 grid. The sections of the dose axis were named “S” (sensitive), “I” (intermediate), and “R” (resilient), while for the time axis, the labels “E” (early), “M” (middle), and “L” (late), were assigned. The final label was then obtained by identifying the earliest and most sensitive point of activation and concatenating the dose and time of the single labels.

### Drug prioritization strategies

#### Dose-dependent SARS-CoV-2 physical interactors

The list of physical interactors with the SARS-CoV-2 were also retrieved from Gordon et al. (Gordon et al., 2020). The list of SARS-CoV-2 physical interactors identified by Gordon et al. was translated to human gene symbols using the R biomaRt package (Durinck et al., 2009). In this manuscript they are referred to as Gordon’s genes. The Gordon’s genes were mapped to the rattus norvegicus ortholog genes using the R biomaRt package (Durinck et al., 2005, 2009). Then, for each drug in the Open TG-GATEs, a score was computed by summing the number of Gordon’s genes that were considered dynamic-dose-responsive by the TinderMIX analysis, and the strength of deregulation as the sum of their log2FC. The log2FC of a dynamic-dose-responsive gene was computed as the mean log2FC of its dynamic-dose-responsive area (Serra et al., 2020a). The drugs were ranked according to this score from the highest to the lowest.

#### Differentially expressed genes in SARS-CoV-2 samples

Similarly, the differentially expressed genes (DEG) identified for the human lung biopsy of SARS-CoV-2 patient, the A549, Calu-3 and NHBE cell lines infected with (SARS-CoV-2) and the A549 cell line infected with SARS-CoV-2 overexpressing ACE2, were mapped to their corresponding rat genes. Then, for each drug in the Open TG-GATEs a score was computed by summing the number of DEG genes in each SARS-CoV-2 condition, that were identified as dynamic-dose-responsive by the TinderMIX analysis, and their strength of deregulation. This resulted in five different ranks of the Open TG-GATEs drugs, where the drugs that strongly deregulate the same DEG of each SARS-CoV-2 condition are at the top of the list.

#### Connectivity mapping

For each one of the SARS-CoV-2 conditions, the genes were ranked from the most up-regulated to the most down-regulated. For each drug of the Open TG-GATEs, the dynamic-dose-responsive genes identified by the TinderMIX analysis were divided into two groups depending on whether their log-fold-changes were monotonically increasing or decreasing in respect to the dose.

The Gene Set Enrichment Analysis (GSEA), based on the Kolmogorov-Smirnov test (Subramanian et al., 2005), was used to compute the pairwise similarity between the Open TG-GATEs drugs and the SARS-CoV-2 conditions. The Kolmogorov-Smirnov test can be used to compare a sample with a reference probability distribution. The Kolmogorov-Smirnov statistic was used without the absolute value in order to preserve the sign (Napolitano et al., 2016). This helps understanding if the increasing and decreasing dynamic-dose-responsive genes derived from the Open TG-GATEs drugs are up- or down-regulated in the SARS-CoV-2 conditions. Thus, for each of the SARS-CoV-2 conditions, and for increasing or decreasing dynamic-dose-responsive behaviour, the Open TG-GATEs drugs were ranked based on their capability to reverse the transcriptomic alterations due to the SARS-CoV-2 infection weighted by the GSEA statistical relevance.

#### Drugs targets and co-expression analysis

All data annotated in the OpenTargets database^5^ (Ochoa et al., 2021) were retrieved as a compressed JSON file. These data contain the drug–targets associations used in this study. The targets were mapped on the previously computed co-expression network and their rank from the INfORM (Marwah et al., 2018) strategies were retrieved. The drugs were ranked according to the median rank of their corresponding targets. In this way, drugs whose targets are more central to the network are ranked at the top of the list.

#### Drug ranking with Borda

Eventually, all the lists of ranked drugs were merged together by using the Borda method (Schimek et al., 2015) implemented in the R package TopKLists. In this way, a final consensus on the drug ranks was identified.

### Relevant chemical substructures

A GSEA analysis was performed to identify the presence of statistically enriched chemical substructure in the drugs ranked at the top of the list. A binary matrix with 881 chemical substructures and 129 drugs was created. A drug was assigned to the list of a specific substructure if that drug contains the substructure. Again, a substructure was considered statistically enriched if the *p* value of the GSEA was lower than 0.05.

### Quantitative Structure–Activity Relationship Analysis

#### Data preprocessing

A quantitative structure–activity relationship (QSAR) analysis (Gramatica, 2020) was performed on the Open TG-GATEs drugs using the pairwise levels of differential expression of a chosen set of genes as the response variable. Namely, *ACE2, TMPRSS2, CTSB* and *CTSL* were used. For each gene, the expression levels at all dose levels and time points were considered simultaneously. The initial dataset for each gene comprised the pairwise differential expression levels at all the doses and all the time points.

In order to differentiate the expression values, a set of dummy variables were added to represent both the dose level and the time points. Substructure fingerprints for the Open TG-GATEs drugs were retrieved from PubChem^6^ (Kim et al., 2016) by querying it by CID identifiers. Each differential expression level was represented by the set of binary fingerprints retrieved by PubChem plus seven indicator variables: three to represent the dose levels, and four to represent the sacrifice periods. Thus, the size of each initial dataset was of 1463 differential expression levels times 888 binary features.

Each of the 881 bits forming the fingerprint indicate the presence or absence of particular chemical substructures, ranging from counts of single atoms, to the presence and the type of bonds between atoms, to more complex substructures like aromatic rings.

Since the fingerprint data matrix was sparse, it was preprocessed by removing all the fingerprints absent in any drug of the dataset manipulating the data matrix with the python module pandas (McKinney, 2010). After this step, the number of variables to use in the modeling phase was further reduced by evaluating the correlation coefficient between each pair, and, to obtain a valid distance metric among the features, the correlation matrix C was transformed according to 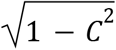 (Van Dongen and Enright, 2012). DBSCAN (Ester et al., 1996) clustering implemented in the python module scikit-learn (Pedregosa et al., 2011), was then applied with parameters epsilon=0.1 and the minimum number of samples as 2. In this way an automatic grouping of the most correlated variables was obtained. Finally, each cluster of correlated variables is compressed into a single one. The resulting data matrix was reduced to about 400 variables.

#### Modeling

After preprocessing, the data were modeled using a gradient boosting machine (GBM) (Friedman, 2001) from the scikit-learn module (Pedregosa et al., 2011), with decision trees as weak learners and the squared loss function. The number of estimators was fixed to 500 for computational constraints and instead we explored different regularization parameters configurations. A grid of possible parameter settings was explored using a 5-fold cross-validation repeated 25 times for each combination of parameters. The grid of parameters was defined as follows,

- subsample, the proportion of samples to use to train each individual tree ranged in [0.2, 0.8],
- max_features, the number of features to evaluate at each node ranged in [1, 50],
- learning_rate, the magnitude of the learning rate varied between [0.001, 0.1],
- max_depth, the maximum height of each decision tree ranged in [1, 5],
- criterion, the function used to evaluate each split was one of {“mse”, “mae”, “friedman_mse”}. Performances of each fitted model were evaluated on the corresponding held-out fold. Due to the lack of an external validation dataset, only internal validation of the fitting was performed. For this reason, the parameters corresponding to the most regularized (i.e. parsimonious) models within one standard deviation from the minimum error achieved were chosen (Hastie et al., 2009).

The best model parameters were identified as subsample=0.5, learning_rate=0.1, max_features=8, criterion=”mse”, max_depth=3 which resulted in a trade-off between validation accuracy, regularization and computing time. The best performing models had a validation RMSE loss of 1.5 ± 0.6.

When the most appropriate parameters were selected for each gene, the best models were fit again using the whole dataset, to obtain the final predictors. Since each drug fingerprint representation appears more than once in the dataset due to the different sacrifice periods and dose levels, care must be taken when considering splits of the dataset into train and validation sets. To this end, every model trained in these experiments was fit and evaluated on datasets split at the drug level, meaning that all instances of a drug are in either the training or validation sets. This is to avoid any information leakage that could happen when some of the dose levels or time points of the same drug are split across the training and validation sets.

#### Chemical substructure relevance

After fitting, each model was exploited to identify the most relevant fingerprints, which, on average, are mostly associated with the over- or under-expression of the analysed genes. To this end, the partial dependence (Friedman, 2001) of the predicted outcome on each fingerprint was computed. Since the optimal selected models fitted decision trees with a maximum depth higher than 1, feature interactions made the ranking slightly unstable, so the fit was repeated for each model 250 times and the relevance of each molecular substructure across the runs was averaged. Finally, each feature was ranked based on its contribution to the average predicted level of differential expression. Each molecular substructure was considered as relevant for under-(resp. over-) expression if the 75th (resp. 25th) percentile of the partial dependence of the response variable is lower (resp. higher) than 0.

### Chemical substructures in screened drugs

A dataset of 6975 chemical compounds screened for activity against several cytotoxicity endpoints from the literature was collected (Ellinger et al., 2020; Gordon et al., 2020; Heiser et al., 2020). For each considered pair of drug and endpoint, an activity threshold was defined in the same way as in the original articles. These activity thresholds were used to define an activity binary variable for each screened drug. The same kind of fingerprints of chemical substructures were also collected for this dataset and a chi^2^ statistical test was performed on each chemical substructure feature against the activity variable to identify the chemical substructures statistically relevant (*p* value < 0.05) to the activity threshold.

### Drugs prioritization

The DrugBank database^7^ v. 5.0 was retrieved for this study. DrugBank (Wishart et al., 2006, 2018) contains 13579 drug entries, including 2635 approved small molecule drugs, and over 6375 experimental drugs. The DrugBank IDs were matched with the PubChem^8^ (Kim et al., 2016) Compound ID (CID) using the PubChem Identifier Exchange Service^9^. For the 8775 matched compounds, substructure fingerprints were retrieved from PubChem by querying it by CID identifiers.

To determine the overall ranking of the DrugBank dataset, we defined the final set of relevant substructures as the union of the fingerprints derived from each single bioinformatics and cheminformatics approach. We further constrained the overall set of relevant substructures by considering its intersection with the set of relevant substructures derived from the screened drugs found in the literature. The drugs were ranked according to the Tanimoto similarity (Bajusz et al., 2015) with respect to the molecular substructures relevant to the models, and considered as possible candidates for further investigation.

### Structural comparison of 7-hydroxystaurosporine and bafetinib

The structures of 7-hydroxystaurosporine and bafetinib (Figure 5) were generated in 3 dimensions and minimized by applying the MM2 energy-minimization method in ChemBio3D (ChemDraw^®^ Professional v20, PerkinElmer Informatics, Inc). The minimized structures were overlaid by steric fields and the similarity was calculated in Discovery Studio Visualizer v21.1 (Dassault Systèmes Biovia Corp).

### Image analysis for syncytia quantification

The fluorescent images (2048 × 2048 pixels) of cells stained with the nuclear dye was used to train a deep convolutional neural network to segment first the nuclei and then the entire cell using the fluorescent signal of the N immunostaining. For the training, we first used 18 images that according to visual inspection faithfully represented the variation (cell size, morphology, average intensity) of the whole dataset. We then randomly extracted a smaller area of 1024 × 1024 sized from each image, and normalized intensities by dividing with the global upper bound of the intensities found in the original dataset and converted to 8-bit format. These selected images were used to manually segment the contour of each N-labelled cell. After this fast initial manual annotation, we utilized image augmentation techniques to prevent the model from overfitting. We created an augmentation pipeline containing 7 main transformations where each transformation instance is applied with a probability of 0.5 implemented using the Numpy scientific programming library and the Pillow package^10^ for Python (Harris et al., 2020). Some of the transformations have input parameters, in this case, the parameter is sampled from an interval divided by step size of 0.01. We constructed 100 augmented instances of each image, therefore we have (100 + 1) × 18 = 1818 images in the extended training set. We then trained the state of the art Mask R-CNN (He et al., 2017) instance segmentation algorithm to detect the cell instances with transfer learning by fine tuning a previous cytoplasm segmentation model to this task. We trained the model through 13 epochs (full model 10 epoch and one epoch for the layers 4+, 3+ and heads), where each epoch contains 2000 steps with batch size of 1.

### Cell culture

HEK-293T-ACE2-TMPRSS2 stable expressing human ACE2 and TMPRSS2 have been previously described (Cantuti-Castelvetri et al., 2020). Caco-2 cells stably expressing human ACE2 were generated by transduction with third generation lentivirus pLenti7.3 ACE2-EGFP, where the expression of EGFP if guided by an internal ribosome entry site downstream of the ACE2 coding sequence in the same messenger RNA. EGFP positive cells were isolated by FACS sorting. All cells were grown in DMEM media supplemented with 10% fetal calf serum (FCS), pen/strep, L-glutamine, and passaged 1:10 (HEK-293T-ACE2-TMPRSS2) or 1:6 (Caco-2-ACE2), every three days.

### SARS-CoV-2 infection

All experiments with wild-type SARS-CoV-2 were performed in BSL3 facilities at the University of Helsinki with appropriate institutional permits. Virus samples were obtained under the Helsinki University Hospital laboratory research permit 30 HUS/32/2018 § 16. The virus was propagated once in Calu-1 cells and once in VeroE6-TMPRSS2 cells before sequencing and storage at −80 °C. Virus stocks were stored in DMEM, 2% FCS, 2 mM L-glutamine, 1× pen/strep as previously described (Cantuti-Castelvetri et al., 2020). Virus titers were determined by plaque assay in VeroE6 TMPRSS2 cells. For testing small molecule inhibitors, cells in DMEM, supplemented with 10% FBS, 1× GlutaMax, 1× pen/strep20, mM HEPES pH 7.2 were seeded in 96-well imaging plates (PerkinElmer cat. no. 6005182) 48 h before treatment at a density of 15,000 cells per well. Drugs, or dimethyl sulfoxide (DMSO) control, were either added 60 min prior to infection, or added 90 min post infection at a multiplicity of infection (MOI) 0.5 plaque forming units per cell. Infections were carried for 20 h in a 37 °C and 5% CO_2_ incubator. Cells were then fixed with 4% paraformaldehyde in PBS for 30 min at room temperature before being processed for immuno-detection of viral N protein, automated fluorescence imaging and image analysis.

### Immunofluorescence, imaging and image analysis

Fixed cells were washed once with Dulbecco PBS containing 0.2% BSA (D-BSA), and permeabilized for 10 min at room temperature in the same buffer containing 0.1% TritonX-100 (W/V, Sigma) and 1 μg/ml Hoechst DNA staining (Thermo Fisher, cat no. H3570). After one wash in D-BSA, cells were incubated for 1 h at room temperature with a 1:2000 dilution of a polyclonal rabbit antibody raised against the viral N protein of SARS-CoV that cross-reacts with the N protein of SARS-CoV (a kind gift of Prof. Ilkka Julkunen, University of Turku, Finland. (Ziegler et al., 2005)). This antibody cross-reacts with SARS-CoV-2 N protein. Following two washes in D-BSA, cells were incubated with a fluorescently labelled goat anti-rabbit antibody (Molecular Probes) at a dilution of 1:1000 for 1 h at room temperature. After two washes in PBS, cells were either stored in the same buffer at 4 °C or imaged directly with a Molecular Device Nano high-content microscope using a 10× objective. To determine the percentage of infected cells, the CellProfiler 3 open source software was used^11^. Nuclei stained with Hoechst were detected using the Otsu algorithm of the CellProfiler3, and infected cells identified based on fluorescence intensity of immunostained N in the perinuclear area of each cell, using a threshold of fluorescence empirically determined such that less than 0.01% of non-infected cells was detected as positive. The relative number of infected cells was calculated by dividing in each well the number of N-positive nuclei by the total number of Hoechst-positive nuclei.

### Quantification of inhibition

Image handling, quantification, and analysis of fluorescence images were performed as described above using Cell profiler 3 or a custom made machine learning algorithm. For each experiment, 9 images were acquired and >2000 cells analyzed. Each experiment was repeated 3-4 times and values indicated in each figure represent the average and standard deviation of all repetitions. Analysis of significance was performed using a one-tailed *t*-test to identify drugs with an infection rate lower than the DMSO. One asterisk= *p*<0.05; two asterisks= *p*<1e-5 after Bonferroni correction.

### Viral infection rate concentration-dependent analysis

A dose-dependent analysis was performed with the strategy implemented in the BMDx tool (Serra et al., 2020b), to test if bafetinib, 7-hydroxystaurosporine, and their combination, reduce the viral infection rate in a dose-dependent manner. For the analysis, the BMR threshold was set to 10% difference with respect to the controls. The linear, power 2, power 3, power 4, exponential, and hill functions were fitted to the data. The optimal fitting model was selected as the one with the lowest Akaike information criteria and used to estimate the BMD, BMDL and BMDU values. The number of replicates used in the concentration-dependent analyses varied from 3 to 15 depending on the concentration level.

## Acknowledgments

R.P. and J.Y.-K. acknowledge the support from the Strategic Research Council at the Academy of Finland (SRC-SUDDEN, Project No. 320210). E.T. and P.H. acknowledge the support from the LENDULET-BIOMAG Grant (2018-342), from the European Regional Development Funds (GINOP-2.3.2-15-2016-00006, GINOP-2.3.2-15-2016-00026, GINOP-2.3.2-15-2016-00037), from the H2020 (ERAPERMED-COMPASS, DiscovAIR), and from the Chan Zuckerberg Initiative (Deep Visual Proteomics). O.V. acknowledges the support from Academy of Finland (grant number 336490), Jane and Aatos Erkko Foundation, EU Horizon 2020 program VEO (874735), and Helsinki University Hospital Funds (TYH2018322). P.A.S.K. acknowledges the support of Orion Research Foundation Sr. G.B. and R.O. acknowledge the support from Academy of Finland (grant number 318434). R.O. also acknowledges the University of Helsinki graduate program In Microbiology and Biotechnology. D.G. acknowledges support from Academy of Finland (grant number 322761), European Union Horizon 2020 Programme (H2020) NanoSolveIT under grant agreement n° 814572, Novo Nordisk Foundation (Grant 0066176), Business Finland (Grant “Nextcast”). The authors are grateful to Troy Faithfull for his critical comments on the manuscript.

## Author Contributions

Conceptualization: D.G. Methodology: A.S., M.F., A.F., R.P., E.T., L.C., P.H. Software: A.S., M.F., A.F., E.T., L.C., P.H. Validation: R.O., G.B. Formal Analysis: A.S., M.F., A.F., R.P., E.T., L.C., P.H., G.B., D.G. Investigation: A.S., M.F., A.F., R.O., R.P., E.T., L.C., G.d.G., P.A.S.K., L.A.S., A.P., V.C., O.V., P.H., A.D.L, J.Y.-K., G.B., D.G. Resources: L.A.S., O.V., G.B., D.G. Data curation: A.S., M.F., A.F., R.O., E.T., L.C., P.H., G.B. Writing - Original Draft: A.S., M.F., A.F., R.P., L.C., G.d.G., P.A.S.K., L.A.S., A.P., D.G. Writing – Review & Editing: A.S., M.F., A.F., R.O., R.P., E.T., L.C., G.d.G., P.A.S.K., L.A.S., A.P., V.C., O.V., P.H., A.D.L., J.Y.-K., G.B., D.G. Visualization: A.S., M.F., R.P., L.A.S., G.B. Supervision: P.H., J.Y.-K., G.B., D.G. Project Administration: G.B., D.G. Funding Acquisition: R.O., E.T., P.H., O.V., P.A.S.K., G.B., D.G.

## Footnotes

1 https://clinicaltrials.gov/ct2/show/NCT04498936

2 https://www.clinicaltrials.gov/ct2/results?cond=&term=UCN-01&cntry=&state=&city=&dist=

3 https://dbarchive.biosciencedbc.jp/en/open-tggates/download.html

4 http://brainarray.mbni.med.umich.edu/

5 https://www.targetvalidation.org/

6 https://pubchem.ncbi.nlm.nih.gov

7 https://www.drugbank.ca

8 https://pubchem.ncbi.nlm.nih.gov

9 https://pubchem.ncbi.nlm.nih.gov/idexchange/idexchange.cgi

10 https://pillow.readthedocs.io/en/3.0.x/reference/ImageEnhance.html

11 www.cellprofiler.org

